# Large-scale genomic rearrangements are a potential explanation for reproductive isolation in the *Pogonomyrmex* dependent-lineage system

**DOI:** 10.64898/2026.05.12.724356

**Authors:** Felix Glinka, Yoann Pellen, Zeev Frenkel, Kimberly K O Walden, Deborah M. Gordon, Eyal Privman

## Abstract

Genetic variation is the raw material for evolution. One source of variation is chromosomal rearrangements, which can bring genes together and form genetic linkage. Rearrangements can also suppress recombination and gene flow, as in the case of sex chromosome evolution. We conducted the first population genomic study of the red harvester ant *Pogonomyrmex barbatus* to investigate genomic rearrangements that differentiate the lineages J1 and J2 in the “dependent-lineage system” (also known as “social hybridogenesis”). In this unusual reproductive system, males and females from different lineages mate to create hybrids, yet these hybrids develop into sterile offspring (workers), and so the two lineages remain reproductively isolated. We sequenced high-quality reference genomes for the two lineages to search for a potential explanation of the suppression of gene flow between them. Comparison of the two genome assemblies revealed multiple large-scale genomic rearrangements, all of which occurred in the J1 lineage. The rearrangements formed some of the largest J1 chromosomes, including the largest scaffold in the assembly that was formed by at least two translocation events and additional intra-chromosomal rearrangements. The translocations brought together 118 odorant receptor (OR) genes on this rearranged chromosome, 44 of which are 9-exon ORs, which are implicated in chemical communication in ants. We also identified an enrichment of transposable elements in a large synteny gap between the translocated segments. The discovery of multiple translocations that formed large rearranged chromosomes provides a potential explanation for the reproductive isolation between the pair of dependent lineages in this system, and opens the way for the study of the molecular genetic basis of an intriguing evolutionary phenomenon in these and in other ant lineages.

## Introduction

Various mutational mechanisms can give rise to chromosomal rearrangements that affect the structure of chromosomes and shape genome evolution (Ranz et al., 2001). Rearrangements can occur through inversion, fission, fusion, or translocation. Such large-scale mutations alter genome organization with potential functional and evolutionary consequences. Chromosomal rearrangements are considered a fundamental mechanism for suppression of recombination and gene flow between populations, potentially leading to speciation (Rieseberg, 2001).

The evolution of sex chromosomes is the best-studied example of the effect of chromosomal rearrangements. Some animal groups, including insects and frogs, display a dynamic evolution and turnover of sex determination mechanisms (Beukeboom and Perrin, 2014; Blackmon & Demuth, 2015; Ezaz et al., 2009; Kaiser & Bachtrog, 2010; Uno et al., 2008). Chromosomal rearrangements are implicated in the evolution of neo-sex chromosomes, having a key role in the suppression of recombination between the X and the Y chromosomes (Ohno, 1967; Beukeboom and Perrin, 2014). Subsequently, genes are translocated to the sex chromosomes and become sex linked (Castillo et al., 2010; Meisel et al., 2020; Traut, 2010). Non-recombining Y chromosomes accumulate repetitive sequences and transposons, and ultimately degenerate over long evolutionary time (Dimitri & Junakovic, 1999; Beukeboom and Perrin, 2014).

Beyond sex chromosomes, so-called “supergenes” evolve by similar mechanisms to code for a wide range of complex polymorphic traits in animals and plants, including butterfly wing patterns and floral morphology (Schwander et al., 2014). Multiple supergenes have been discovered in ants. The first was described in the fire ant *Solenopsis invicta*, in which a “Y-like social chromosome” harbors a supergene that is associated with queen number (Wang et al., 2013). Three inversions in the Y-like chromosome can explain the suppression of recombination in this supergene (Yan et al., 2020).

While an inversion is an intra-chromosomal rearrangement that suppresses recombination in a limited chromosomal region, inter-chromosomal rearrangements (fusions, fissions, and translocations) can lead to complete failure of meiosis and thus to sterility. The best-known example is that of the mule, which receives a different number of chromosomes from its horse mother (2n=64) and donkey father (2n=62) (Benirschke et al., 1962). It is not clear whether rearrangements are the primary mechanism for reproductive barriers, but they are widely considered to have an important role in the evolution of reproductive isolation that ultimately leads to speciation (Rieseberg, 2001). Chromosomal rearrangements can also create reproductive barriers between sympatric populations that undergo local adaptation to different ecological niches (Fishman et al., 2013).

Here, we investigate chromosomal rearrangements as a potential explanation for reproductive isolation between a pair of “dependent lineages” in a population of *Pogonomyrmex barbatus* harvester ants that evolved an unusual reproductive strategy known as a “dependent-lineage system” or “social hybridogenesis” (Volny & Gordon, 2002; Julian et al., 2002; Helms Cahan & Keller, 2003). Multiple dependent lineage pairs were discovered in the *Pogonomyrmex barbatus*/*rugosus* species complex, including F1/F2, G1/G2, H1/H2, and J1/J2, alongside other closely related populations that do not have dependent lineages (Schwander et al., 2007). The extensive distribution of these dependent-lineage populations was documented throughout Southern USA and Northern Mexico (Anderson et al., 2006; Schwander et al., 2007; Mott et al., 2015).

In these dependent-lineage populations, matings between a queen and a male of the same lineage produce reproductive females, while matings between a queen and a male of different lineages produce workers, which are sterile. Thus, the lineages are effectively evolving as two reproductively isolated populations. This appears to be at least in part the result of a genetic caste determination mechanism, whereby a female embryo evolves into either a queen or a worker depending on some genetic factor (Helms Cahan et al., 2002; Volny & Gordon, 2002). This is in contrast to the more common environmental caste determination, such as the differential feeding of honeybee larvae with royal jelly that determines queen vs. worker development (Weaver, 1955). Some studies have reported exceptions to genetic caste determination in *P. barbatus*, where 3 out of 80 gynes (virgin queens) were found to be inter-lineage hybrids (Helms Cahan & Keller, 2003; Sirviö et al., 2011). The reproductive success of colonies with a hybrid queen is unknown, but these reports raise questions such as how the development of reproductive hybrids is suppressed genetically or otherwise, and what part this mechanism plays in the evolution and maintenance of the dependent-lineage system over long evolutionary timescales.

The peculiar nature of the dependent-lineage system attracted attention over the past two decades from many researchers who explored potential explanations for its evolution. Many studies have investigated the phylogenetics and population genetics of dependent-lineage systems using mitochondrial and nuclear loci. Helms Cahan and Keller (2003) suggested that the origin of the J1/J2 dependent-lineage system was hybridization between *P. barbatus* and *P. rugosus;* and Helms Cahan and colleagues (2022) suggested a similar origin for other lineage pairs. Anderson and colleagues (2006) argued that the dependent-lineage system first evolved in *P. barbatus,* not through hybridization between species, and then introgressed into *P. rugosus*. Schwander and colleagues (2007) found a ratio of 75/25% *P. rugosus*/*P. barbatus* genetic ancestry in J1, while for J2 has pure *P. barbatus* ancestry. Altogether, previous analyses revealed an extensive history of hybridization and introgression events among the populations and lineages in this clade. Helms Cahan and colleagues (2023) suggested that hybrid superiority drove the evolution of dependent-lineage systems, possibly related to adaptation to climatic conditions (especially precipitation and seasonality) and resource availability.

Dependent-lineage systems are maintained by the suppression of interlineage gene flow, as interlineage matings produce workers (Anderson et al., 2008). Helms Cahan and Keller (2003) hypothesized that hybrid incompatibility results in worker development, while same-lineage matings can produce fertile females (i.e. queens). Linksvayer (2006) suggested that intra-lineage mating produces female offspring with coevolved cyto-nuclear gene complexes, which develop into fertile females, while inter-lineage mating produces females with disrupted cyto-nuclear complexes (i.e., cyto-nuclear incompatibility), which can develop only into sterile females. The two explanations - incompatible nuclear alleles and cyto-nuclear incompatibility - are not mutually exclusive.

Similar dependent-lineage systems have been reported in other ant species. Most interestingly, the *Messor* genus evolved granivory (seed foraging) and a dependent-lineage system similarly and independently of *Pogonomyrmex*, indicating convergent evolution in the two genera (Romiguier et al., 2016, 2017). Romiguier and colleagues (2017) suggested two ecological explanations for the convergent evolution of dependent-lineage systems in these species: large synchronous mating flights and a granivorous diet. Interestingly, *Messor barbarus* workers can produce hybrid males, which may result in gene flow between lineages (Romiguier et al., 2017). However, it is not known whether such hybrid *Messor* males or the hybrid *Pogonomyrmex* gynes are fertile and can produce viable offspring.

Here, we examine genomic differences between the J1/J2 lineages to investigate whether chromosomal rearrangements are responsible for the reproductive barrier. We produced the first genomic sequence data for the J1/J2 dependent-lineage system. We constructed high-quality genome assemblies for the two lineages, which revealed multiple large-scale genomic rearrangements that created large chromosomes in J1, most notably scaffold 1 of the J1 genome assembly. We then used population genomic data to show that these rearrangements are fixed or nearly fixed in the population, including whole genome sequencing of 12 males from each lineage and reduced representation genomic sequencing of 1101 worker samples. We studied the gene content of the largest rearranged J1 chromosome and found that it is enriched with odorant receptor genes, which are implicated in chemical communication in ants. We propose that this rearranged J1 chromosome contributed to the evolution of the dependent-lineage system and that it may be responsible for the reproductive barrier between the two lineages.

## Results

### Large-scale rearrangements in the J1 lineage

To detect large-scale genomic rearrangements, we sequenced and assembled two high-quality reference genomes from one J1 male and one J2 male (scaffold N50 of 10,579,937 and 10,965,113 bp, respectively). We obtained a telomere-to-telomere assembly for 4 chromosomes in each assembly, and among the largest scaffolds there are 12 J2 scaffolds and 13 J1 scaffolds that have a telomere at one end (Supplementary Tables 1 and 2). We also generated large-scale population genomic sequencing data, including 12 males from each lineage (whole genome sequencing) and 1101 workers from 407 colonies (reduced representation genomic sequencing). The population genomic data are consistent with previous evidence for reproductive isolation between lineages (e.g., Sirviö et al., 2011). We found a value of 0.54 average genome-wide *F_ST_* between the J1 and J2 males, including 13 scaffolds with nearly fixed polymorphic sites (*F_ST_* > 0.9). We observed a U-shaped distribution of heterozygosity in the workers (Supplementary Figure 1), as expected for F1 hybrids between two diverged populations.

The comparison of the genome assemblies from J1 and J2 reveals multiple large chromosomal rearrangements. Figures 1A&B show the homology of the J2 and J1 scaffolds to the genome of *Pogonomyrmex californicus*, which was assembled to chromosome level. Figure 1C plots the homolog of J1 and J2 scaffolds. While the large J2 scaffolds are almost perfectly co-linear with *P. californicus* chromosomes, the J1 alignment shows multiple large-scale rearrangements. This indicates that the J1 genome has undergone extensive rearrangement, whereas the J2 lineage kept the ancestral genome structure. The largest three scaffolds (1, 2, and 3) in the J1 genome are homologous to distinct fragments of multiple large *P. californicus* chromosomes. These results suggest that multiple inter-chromosomal translocations formed the largest chromosomes in the J1 lineage.

**Figure 1:**
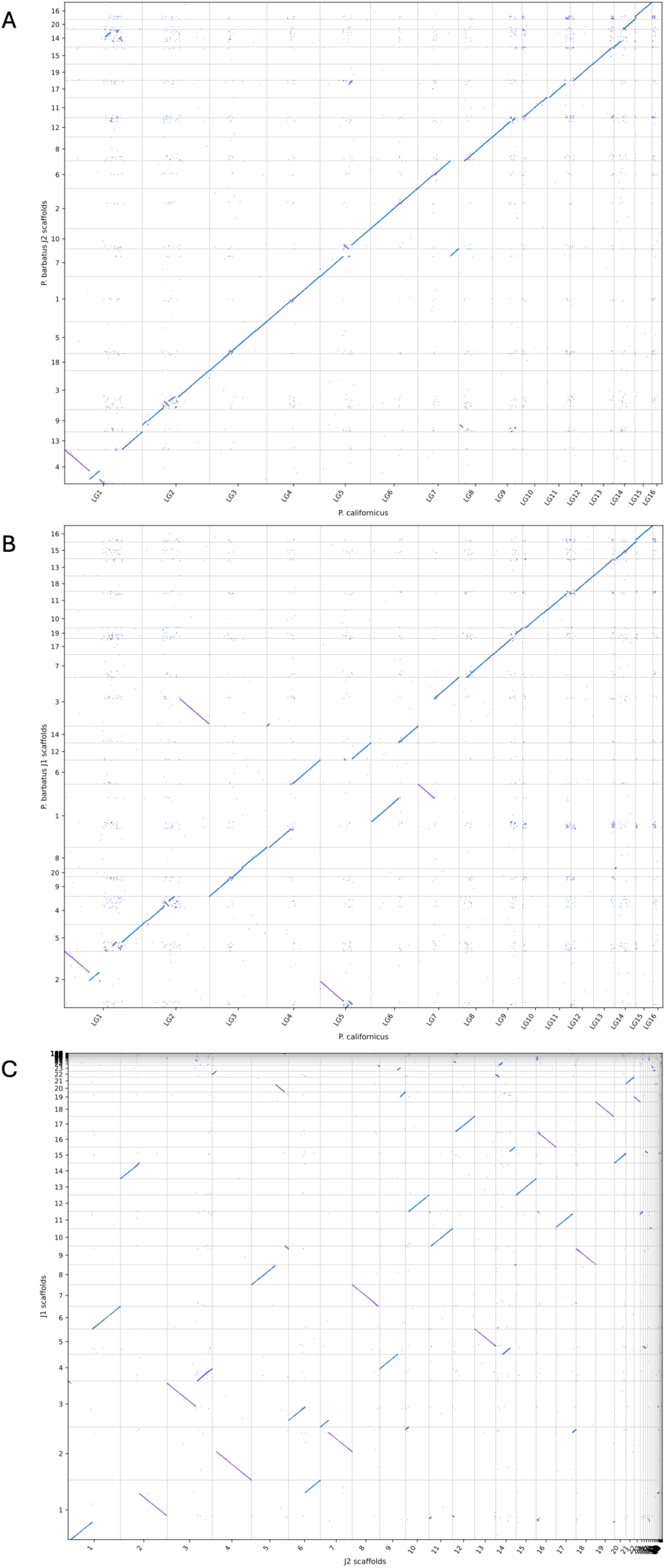
Synteny dot plots of the *Pogonomyrmex barbatus* J1 and J2 genomes and the *P. californicus* genome. Plots are based on genome alignments, and each point represents a segment of sequence homology. **A:** J2 vs. *P. californicus*, **B:** J1 vs. *P. californicus*, **C:** J1 vs. J2. LG corresponds to Linkage Group in *P. californicus*; numbering on the J1 and J2 genomes corresponds to genomic scaffolds. Blue: plus strand, violet: minus strand.

One-to-one orthologous protein-coding genes between the *P. californicus* and the J1 and J2 genomes are plotted in Figure 2 to visualize large syntenic blocks and the large-scale chromosomal rearrangements. We focus on the three largest scaffolds of the P. barbatus J1 genome, which were rearranged relative to the ancestral genome (The complete plot with all chromosomes can be found in Supplementary Figure 2). Most notably, J1 scaffold 1 (29,218,669 bp long) is composed of three large segments that are homologous to segments of J2 scaffolds 1, 2, and 6 (the homologous fragments are 8,498,785; 10,815,236; and 6,368,357 bp long, respectively). J2 scaffolds 1, 2, and 6 are homologous to *P. californicus* linkage groups 4, 6, and 7, respectively, with no apparent large-scale rearrangements. The breakpoints between the large segments in J1 scaffold 1 correspond to breakpoints in the middle of the J2 scaffolds and the *P. californicus* chromosomes, demonstrating that these are indeed translocations and not a result of incomplete assembly of the J2 chromosomes. Note also that J1 scaffold 3 and J2 scaffold 2 are telomere-to-telomere assemblies, indicating that these are fully assembled chromosomes. In combination, these results provide strong support for at least five interchromosomal translocations that rearranged large chromosomal fragments to form at least three of the largest chromosomes in the J1 genome, while there is no evidence for such large-scale rearrangements in the J2 genome.

**Figure 2:**
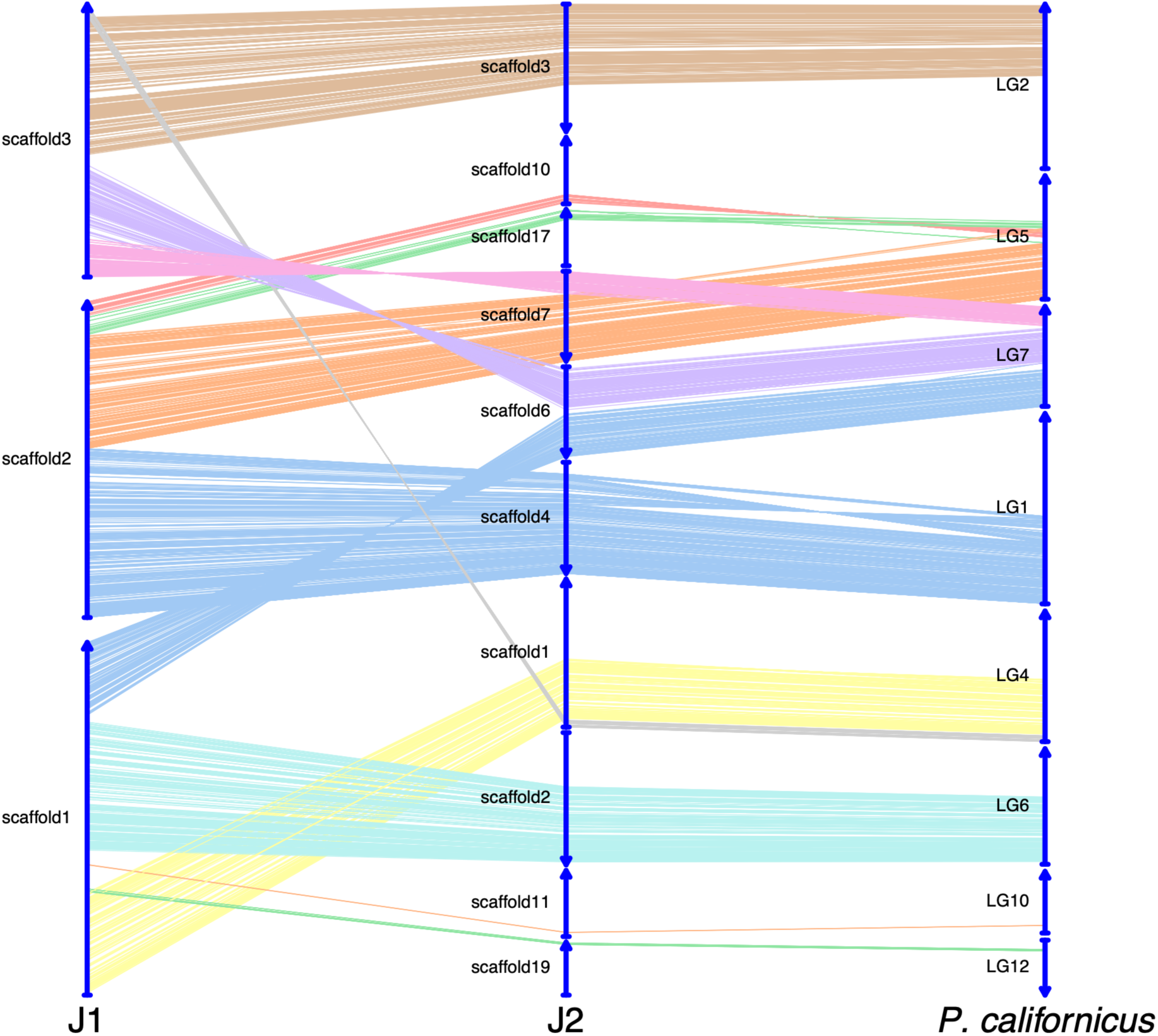
Conservation of synteny illustrated using a river plot comparing the J1 and J2 of *P. barbatus* and *P. californicus* for the three largest scaffolds that were formed in the J1 lineage by rearrangements. Each line represents one-to-one orthologous protein-coding genes.

The synteny plots of J1 scaffold 1 against the other genomes show a large gap without collinear homology (Figures 1 and 2). These results show that this region in J1 scaffold 1 has no homologous chromosome with conserved synteny in J2 and the *P. californicus* genomes. On a closer inspection of J1 scaffold 1 (Figure 3A), the large gap (positions 8,505,971-11,882,941) becomes visible between the alignments to segments of J2 scaffolds 1 and 2. Zooming closer into this region (Figure 3B) reveals that this is a highly rearranged region made up of short fragments of multiple J2 scaffolds (19, 16, 11, and 12) as well as many repetitive sequences.

**Figure 3:**
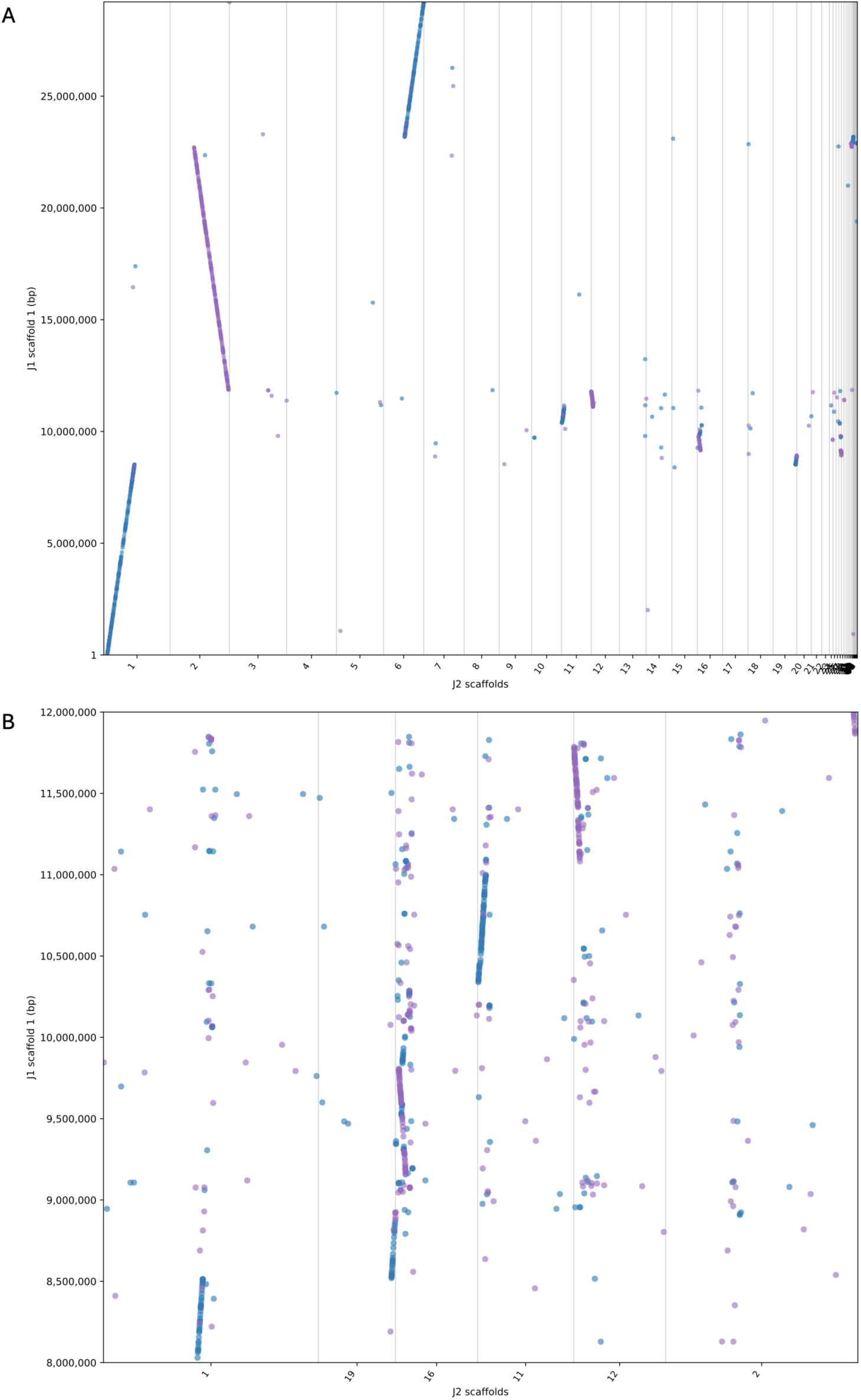
Synteny plots for J1 scaffold 1. **A:** scaffold 1 of J1 vs the whole J2 genome; **B:** Zoom-in on J1 scaffold 1 positions 8,000,000-12,000,000, showing only J2 scaffolds with substantial homology. Violet: plus strand, blue: minus strand.

We confirmed the J1 rearrangements using population genomic data, which show that they are widespread across the population of J1 genomes. We generated whole-genome sequencing data from 12 males from each lineage and surveyed the coverage along each of the two reference genomes for sequencing reads from each of the samples. Generally, high uniform coverage is visible across most of J1 scaffold 1 for all 24 samples (Figure 4A). The 3.3 Mbp synteny gap around position 10 Mbp in the scaffold (Figure 3) shows a dramatic drop in coverage in the J2 samples relative to the J1 samples (Figure 4A). We noted a general trend across the genome: the J2 samples have slightly lower coverage than the J1 samples (see for example J1 scaffold 7 in Supplementary Figure 3A). We suspected that this was due to reference bias, and indeed we observed the reverse pattern in the coverage of the J2 reference genome (see the homologous J2 scaffold 8 in Supplementary Figure 3B). Similarly, J1 samples have an overall higher average coverage than J2 samples for regions of J1 scaffold 1 outside the gap (Figure 4A), and the homologous regions in J2 scaffolds 1, 2, and 6 show a higher average coverage for J2 samples than J1 samples (Figure 4B). Nevertheless, both J1 and J2 samples show uniform coverage for these regions in which synteny is conserved between the lineages.

**Figure 4:**
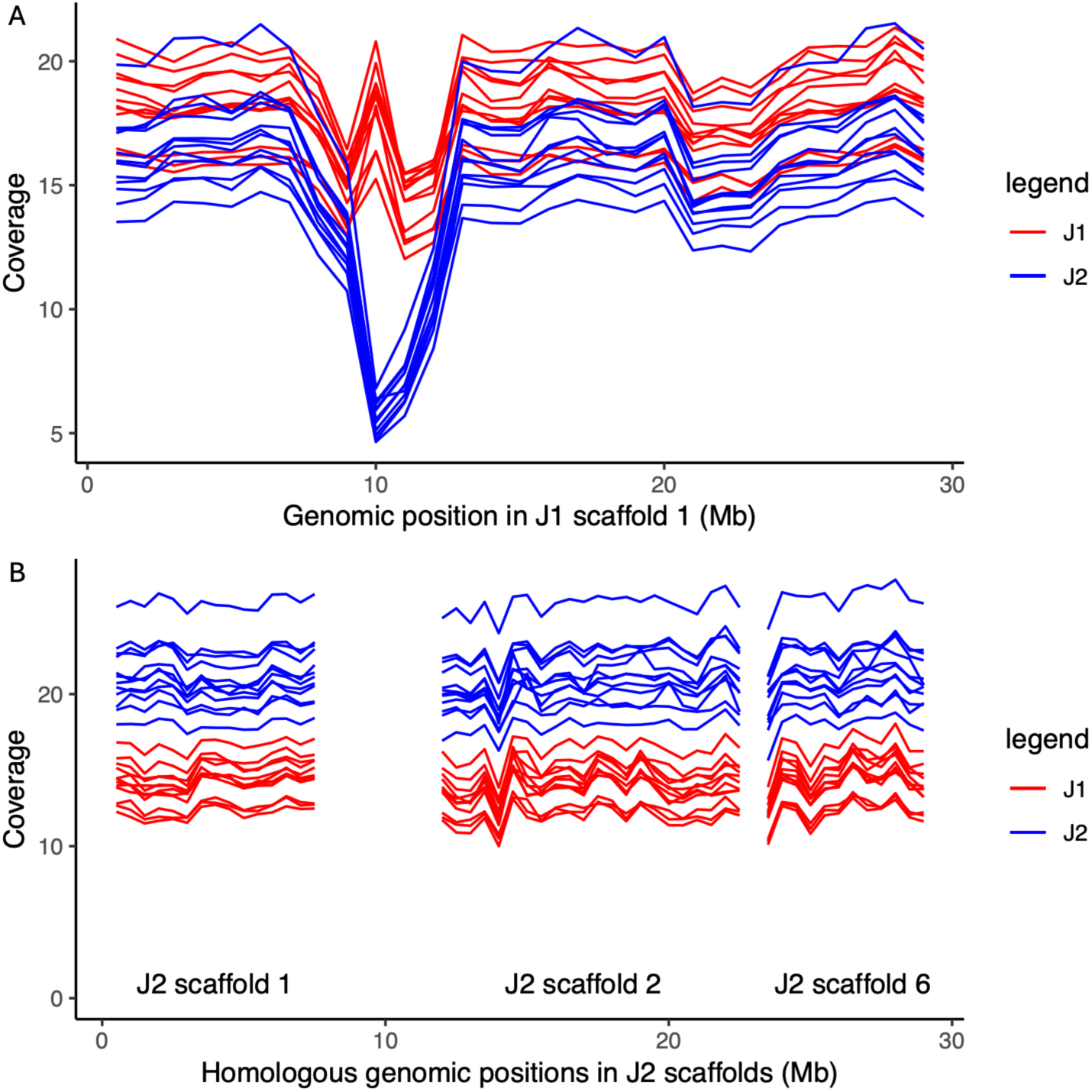
Sequencing coverage of the rearranged J1 scaffold 1 (A) and the homologous regions of J2 scaffolds for J1 (red) and J2 (blue) male samples. Plots show moving average for a window size of 1Mbp.

We then used reduced representation genomic sequencing data from worker samples to confirm the presence of the synteny gap across the population. Workers are inter-lineage hybrids and are thus expected to have sequencing reads only from their J1 genome mapping to the gap region. This would result in low heterozygosity relative to the rest of the genome. The plot of worker heterozygosity along the scaffold is consistent with this prediction (Figure 5). Worker samples have very low heterozygosity in this region, while the rest of the scaffold shows the typical, genome-wide, U-shaped distribution of heterozygosity (Supplementary Figure 1).

**Figure 5:**
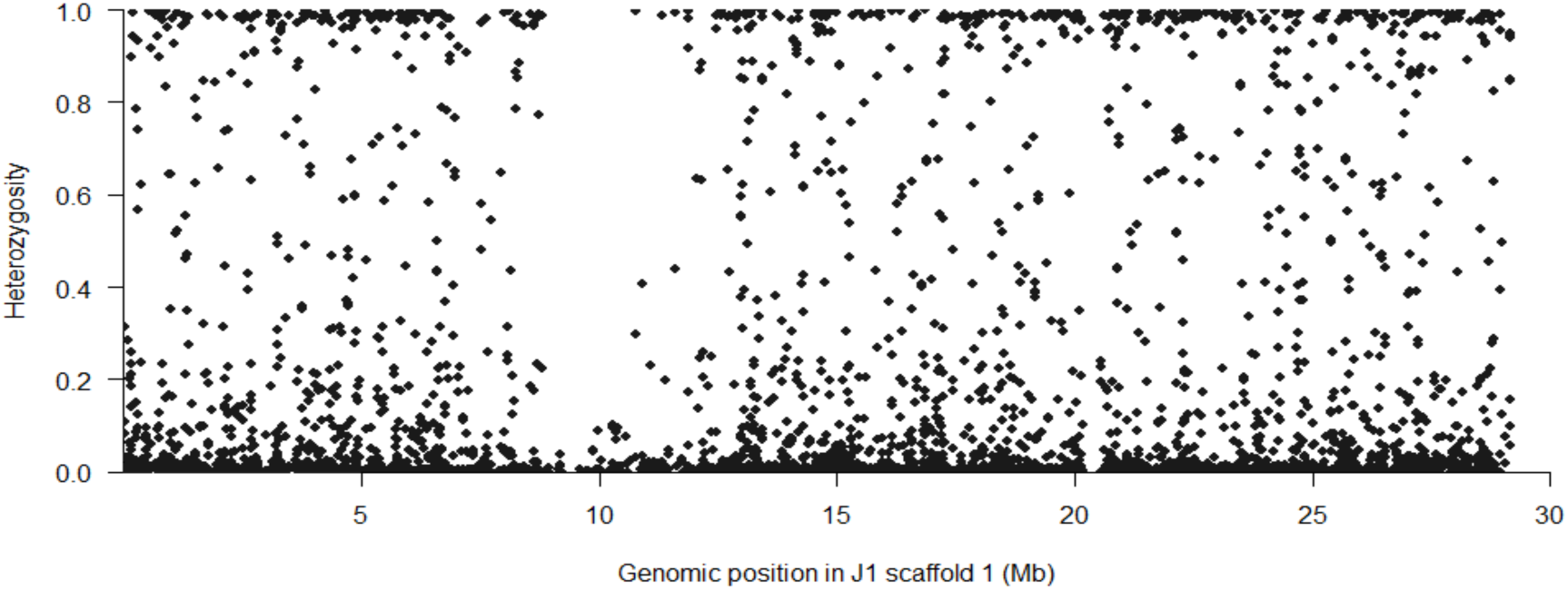
Heterozygosity of SNPs in worker samples along J1 scaffold 1.

### Gene content of the largest rearranged J1 scaffold

We annotated genes in the J1 genome and investigated J1 scaffold 1 to identify gene functions that were brought together in this large chromosome by the translocations. Supplementary Table 4 lists all annotated protein-coding genes in the scaffold.

We found that the rearrangements brought together large odorant receptor (OR) arrays into one large chromosome. The scaffold was strikingly enriched with OR genes (Fisher’s exact test corrected *p* = 4E-24). The scaffold contains 118 of the total of 352 OR genes in the J1 genome. No other functions showed statistically significant enrichment in scaffold 1 (Table 1). Eight out of 25 subfamilies of ORs are represented in the scaffold (Supplementary Table 3), including most notably, 51 out of the 151 genes of the 9-exon subfamily (Fisher’s exact test *p* = 1E-14) and all of the 43 ORs of subfamily L (*p* = 3.99E-43).

**Table 1:** Functional enrichment in genes in J1 scaffold 1. Each GO term was tested for enrichment, and the *p-*values were corrected for multiple testing.

| GO.ID | GO term | Whole genome | Scaffold 1 | Expected | $p$ value | FDR corrected $p$ value |
| --- | --- | --- | --- | --- | --- | --- |
| **Biological Processes** |  |  |  |  |  |  |
| GO:0007608 | sensory perception of smell | 374 | 119 | 45.52 | 2.50E-25 | 2.89E-21 |
| GO:0006183 | GTP biosynthetic process | 7 | 4 | 0.85 | 0.0056 | 1 |
| GO:0009408 | response to heat | 42 | 9 | 5.11 | 0.0076 | 1 |
| **Cellular Component** |  |  |  |  |  |  |
| GO:0016020 | membrane | 3112 | 415 | 371.01 | 6.40E-14 | 1.03E-10 |
| GO:0035578 | azurophil granule lumen | 7 | 4 | 0.83 | 0.0052 | 1 |
| GO:0062129 | chitin-based extracellular matrix | 12 | 5 | 1.43 | 0.0092 | 1 |
| **Molecular Function** |  |  |  |  |  |  |
| GO:0005549 | odorant binding | 356 | 119 | 43.25 | 1.20E-27 | 4.16E-24 |
| GO:0004984 | olfactory receptor activity | 354 | 118 | 43.01 | 2.40E-27 | 4.16E-24 |
| GO:1904315 | transmitter-gated monoatomic ion channel activity involved in regulation of postsynaptic membrane potential | 29 | 8 | 3.52 | 0.0051 | 1 |
| GO:0008010 | structural constituent of chitin-based larval cuticle | 12 | 5 | 1.46 | 0.0099 | 1 |

**Table 2:** Counts of transposon-associated keywords in gene descriptions within the synteny gap in J1 scaffold 1.

| Keyword | Gene count |
| --- | --- |
| transposon | 78 |
| transposase | 43 |
| transposable | 10 |
| mobile element | 1 |
| Gag-Pol polyprotein | 9 |
| virus | 64 |
| SETMAR | 2 |
| Copia | 2 |
| Capsid | 3 |

The scaffold contains seven tandem arrays of OR genes (Figure 6). All 43 L subfamily ORs are in a single tandem array in the third segment of the scaffold, which is homologous to J2 scaffold 6. The 9-exon ORs are in two tandem arrays, one array of 7 ORs in the first segment (homologous to J2 scaffold 1) and an array of 44 ORs in the second segment (homologous to J2 scaffold 2). Thus, the two translocations brought together three chromosomal segments that each contain substantial numbers of ORs to form a large chromosome that is enriched with ORs relative to the rest of the genome.

**Figure 6:**
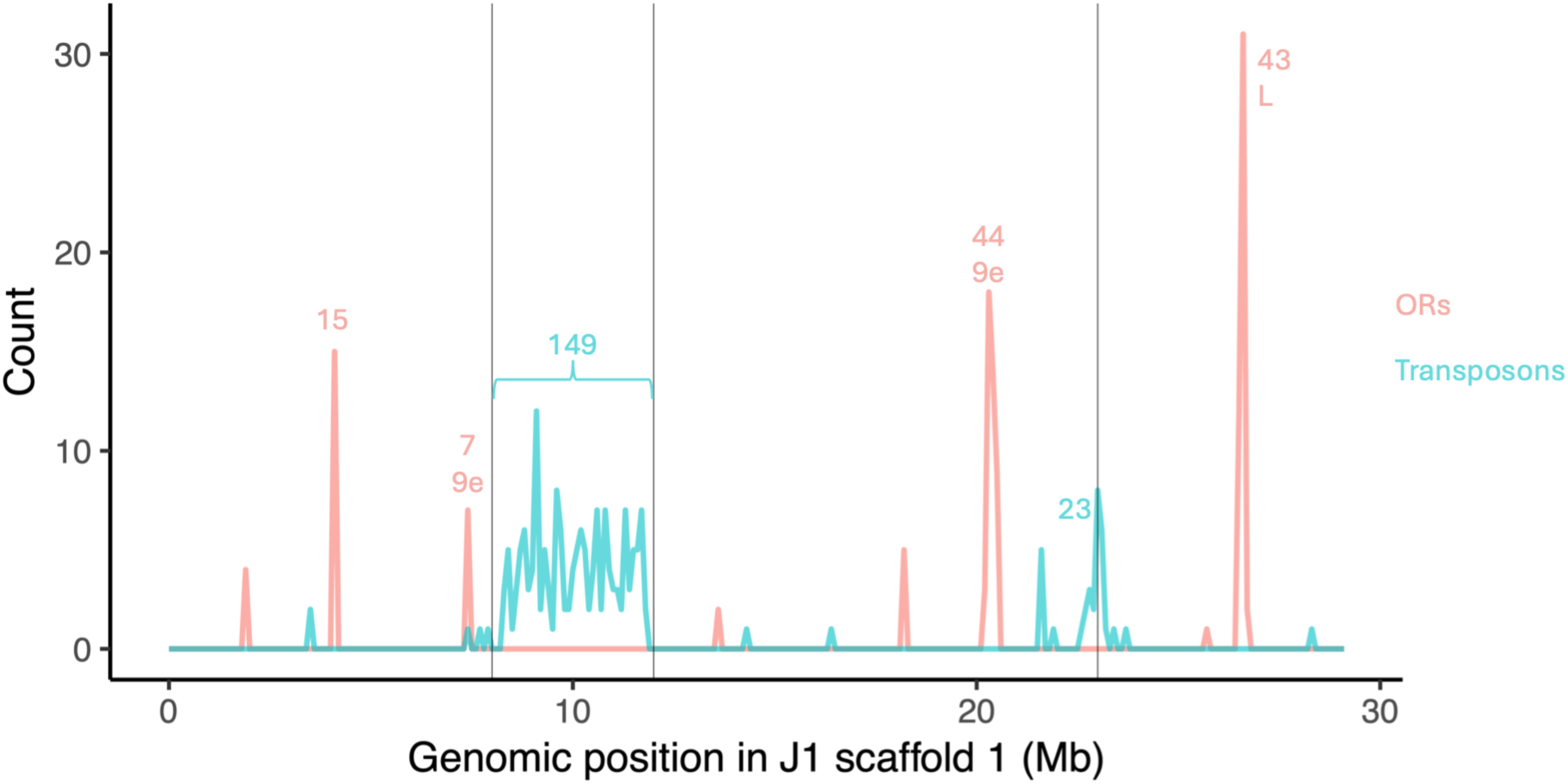
Distribution of odorant receptor gene (OR, red) and genes associated with transposons (cyan) on J1 scaffold 1. Genes are counted in bins of 100,000 bp. Numbers above peaks show the cumulative number of genes in bins. Black vertical lines show the rearrangement breakpoints. OR subfamilies L and 9e (“9-exon”) are highlighted.

Interestingly, a gene tree constructed for the L subfamily OR from eight representative species from diverged ant subfamilies shows that most of the 43 *P. barbatus* ORs are the result of ancient gene duplications in an ancestor of most extant ant species (Supplementary Figure 4). This result implies that this large gene array was formed in an ancient ancestor and remained intact over more than 100 million years of evolution.

Another gene family that was implicated in olfactory response to chemical signaling in ants is that of the odorant binding proteins (OBPs) or pheromone binding proteins. J1 scaffold 1 contains a tandem array of four OBPs in the first segment of the scaffold, around position 1.9 Mbp. Interestingly, these four paralogs are highly similar to each other, suggesting recent gene duplication in the *Pogonomyrmex* lineage. Their closest orthologs in the fire ant *Solenopsis invicta* are *SiOBP3* and *SiOBP4*. *SiOBP3* is the first OBP discovered in the fire ant social chromosome (originally named *Gp-9*) and implicated in the differences between the two social forms in polymorphic *S. invicta* populations (Gotzek et al., 2011; Keller & Ross, 1998; Krieger & Ross, 2002).

In the synteny gap, we found many genes associated with transposons. A total of 149 genes in this region had keywords associated with transposons in their gene description (Table 2), which corresponds to statistically significant enrichment (Fisher’s exact test *p* = 4E-69). The second break point region (between the second and third segments, at position 22 Mbp) contains 23 transposon-associated genes (Figure 6).

## Discussion

Near-chromosome level assemblies of two representative genomes from the J1 and J2 lineages revealed multiple translocations involving large chromosomal segments that created some of the largest chromosomes in the J1 genome. Large-scale population genomic data support the identified rearrangements in the two representative genomes. Thus, the rearrangements in J1 appear to be fixed or nearly-fixed across the population of J1 genomes.

Our genome assemblies achieved high contiguity thanks to a very successful combination of long-read Pacbio sequencing and linked-read Tell-seq data. We also attempted Hi-C but this protocol failed due to some unknown cause in the *P. barbatus* samples (we attempted both male and worker samples). Nevertheless, the high contiguity of the two assemblies (N50 > 10 Mbp; 7 and 9 scaffolds >10 Mbp in J1 and J2, respectively) allowed the confident identification of multiple large scale rearrangements. The comparison between the J1 and J2 genomes and the chromosome-level assembly of the *P. californicus* genome shows that the rearrangements happened in the J1 lineage, while the ancestral chromosomes are conserved in J2. A recent study dated the split between the *P. californicus* and *P. barbatus* lineages to about 30 million years ago based on a phylogenomic analysis of ultraconserved elements (Graber et al., 2024). The conservation of synteny in the J2 genome suggests that chromosome structure was largely conserved in the *P. barbatus* lineage, except in the J1 lineage.

Since this is the first such genomic study of a dependent-lineage system in *Pogonomyrmex*, we do not know whether similar evolutionary dynamics shaped the genomes of lineages of other such systems (F1/F2, G1/G2, H1/H2). A recent study of the dependent-lineage system in *Cataglyphis hispanica* showed that the lineages differ in the number of chromosomes due to a fission or fusion event (Darras et al., 2022). However, this difference is not fixed between the two lineages, and so the authors conclude that it was not a causal mutation for reproductive isolation in this system. Our large population genomic sample shows that the translocations are fixed or nearly fixed in our study population. Thus, they can prevent recombination and lead to suppression of gene flow between the two lineages. It will be interesting to see whether our results are generalizable in future studies of more dependent-lineage populations in *Pogonomyrmex* and other ant lineages such as the *Messor* genus.

The largest rearranged J1 chromosome is made up of three large chromosomal segments, with a 3.3 Mb synteny gap enriched with transposons and other repetitive sequences. Such dramatic rearrangements could prevent recombination between the J1 chromosome and the homologous chromosomes in the J2 genome.

The evolution of the rearranged J1 chromosome can be seen as analogous to the evolution of sex chromosomes, and more generally to the evolution of supergenes. Genomic rearrangements are the key mechanism leading to suppression of recombination between haplotypes in the evolution of sex chromosomes and supergenes (Charlesworth, 2017). While some studies identified translocations involved in the formation of sex chromosomes in fish, frogs, and lizards (Ezaz et al., 2009; Pennell et al., 2015; Uno et al., 2008), most of the rearrangements in sex chromosomes and supergenes are intra-chromosomal inversions, for example, in the evolution of an incipient sex chromosome in papaya (Wang et al., 2012) and in the evolution of a supergene associated with social polymorphism in fire ants (Yan et al., 2020).

In sex chromosomes and supergenes, the suppression of gene flow is limited to the non-recombining region. By contrast, the rearrangements we discovered may result in a failure of meiosis in inter-lineage hybrid female offspring, which would render them sterile. Production of sterile gynes is a substantial waste of resources and could have a fitness cost for the colony. Such a fitness cost may lead to selection for the evolution of the dependent-lineage system, in which only same-lineage matings develop into reproductive females. There are reports of rare hybrid gynes (Sirviö et al., 2011) and Helms Cahan and colleagues (2022) suggested that there may be interlineage gene flow via the male offspring of workers produced in colonies that have lost the queen. However, there is no evidence that such hybrid gynes or males are fertile. The rarity of such hybrid reproductive offspring can be explained by the fitness cost of sterility due to the genomic rearrangements. We propose that the fitness cost of the production of sterile gynes resulted in the evolution of genetic caste determination so that almost no hybrid gynes are produced.

The translocations in J1 formed a large chromosome that is enriched with olfactory genes (Table 1). A striking number of 118 out of the total of 352 OR genes were brought together on this chromosome, including the entire L subfamily of ORs and 51 of the 9-exon ORs. The OR gene family is significantly expanded in ants, which was suggested to be due to the evolution of elaborate chemical communication (Smith et al., 2011a; Smith et al., 2011b). The large numbers of ORs that were translocated to the same chromosome implicate this rearranged chromosome in chemical communication, especially because of the presence of 51 9-exon ORs. The 9-exon is the most expanded OR subfamily in ants, which were shown to respond to cuticular hydrocarbons (McKenzie et al., 2016; Pask et al., 2017; Slone et al., 2017). Cuticular hydrocarbons (CHCs) are a major group of signaling compounds used for chemical communication by insects and especially in social insects (Blomquist & Bagnères, 2010). Thus, this rearranged J1-specific chromosome may code for chemical signalling and recognition functions that facilitate the functioning of the dependent-lineage system.

Differences were reported in the CHC profiles of males from the different lineages (Volny et al., 2006). Olfactory detection of these chemical signals by ORs on the rearranged chromosome may allow gynes to determine if they have mated with males of both lineages, which is vital for the establishment of a fully functioning colony. The rearranged chromosome ORs may also be involved in discrimination between worker- and gyne-destined larvae (Helms Cahan et al., 2002). Similar discrimination powers of *P. barbatus* are known in the context of task recognition based on CHC profiles of workers (Greene & Gordon, 2003). In the fire ant *Solenopsis invicta*, unsaturated CHCs allow workers to discriminate between monogyne and polygyne queens (Eliyahu et al., 2011; Zeng et al., 2021). Due to genetic caste determination in *P. barbatus* colonies, worker- and gyne-destined eggs are produced in the ratio corresponding to the queen’s mates. However, gynes do not emerge from colonies before they are five years old and the colony has reached mature size (Gordon 1995). Thus, novel olfactory functions in dependent-lineage populations may allow nurse workers to distinguish worker- vs. gyne-destined eggs or larvae and eliminate the latter. An alternative explanation would involve discriminating hybrid vs. pure lineage larvae, which facilitates their differential feeding by the nurses to induce development into workers or gynes, respectively. Such potential mechanisms were proposed and discussed at length in previous publications (e.g., Helms Cahan et al., 2002; Sirviö et al., 2011).

Transposons and other repetitive sequences accumulate in Y chromosomes because they never combine (Dimitri & Junakovic, 1999; Steinemann & Steinemann, 2005; Beukeboom and Perrin, 2014). Similar evolutionary dynamics also characterize Y-like supergenes, such as the chromosome that is associated with the polygyne form in fire ants (Fontana et al., 2020). The large synteny gap in J1 scaffold 1 is highly enriched with genes associated with transposons (Figure 6). Transposons are also associated with the second breakpoint in this chromosome (around position 22 Mb; Figure 3). This suggests that the J1 rearrangements created chromosomal regions with suppressed recombination.

Further studies are needed for a comprehensive understanding of the genomic basis of the origin and evolution of the dependent-lineage system. Genomic sequencing of additional populations and phylogenomic analysis are required to fully resolve the history of genome evolution in this *Pogonomyrmex* species complex. Such studies could shed light on the role of hybridization between *P. barbatus* and *P. rugosus* in the evolution of the dependent-lineage system - whether hybridization was the origin of the system (Helms Cahan and Keller, 2003) or whether the dependent-lineage system first evolved first in *P. barbatus* and later introgressed into *P. rugosus* (Anderson et al., 2006). Phylogenomic analysis could determine whether the rearrangements we discovered in the J1/J2 population evolved in an ancestor of multiple dependent-lineage pairs (including the F, G, and H lineages) or whether they are specific to the J1/J2 system. If the rearrangements evolved recently in the J1/J2 system, it would be interesting to see whether analogous rearrangements occurred in the other *Pogonomyrmex* dependent-lineage systems. That is, whether rearrangements are the shared mechanism of convergent evolution of such reproductive systems. The same question applies also to other more distant dependent-lineage systems, such as in the *Messor* genus.

## Materials and Methods

### Samples

We collected samples of *Pogonomyrmex barbatus* from the long-term study site of the Gordon lab near Rodeo, New Mexico, US (Sundaram et al., 2022). Worker samples that were collected between 2001 and 2023 from the field site were used for sequencing, including 1101 worker samples from 407 (for details please see Glinka et al., 2026). Male samples were collected from mating swarms in the summer of 2024. This population was studied extensively and multiple previous studies conducted genetic analyses that are in line with the expectations under a dependent-lineage system of colony reproduction (e.g. Volny & Gordon 2002; Volny et al. 2006).

### Sequencing of a reference genome from each lineage

DNA was extracted from the whole body of each male sample using the Qiagen DNeasy blood and tissue protocol followed by ethanol precipitation. We identified the lineage of each sample based on their mitochondrial haplotypes. We used the universal insect *cox*1 primers modified for *P. barbatus* to amplify and sequence the haplotype. We then constructed a phylogenetic tree of the haplotypes to assign each sample to the J1 or the J2 clade, which were identified based on previously published sequences (Helms Cahan & Keller, 2003). To obtain high-quality genome references, one male of each lineage was sequenced and assembled based on a combination of long-read PacBio HiFi sequencing and the linked-read Tell-seq protocol. PacBio sequencing coverage was on average 41X for J1 and 30X for J2, with an average read length of 7 Kbp. The Tell-seq average coverage was 459X for J1 and 402X for J2. First, we assembled the long reads with *Hifiasm* (Cheng et al., 2021). Assembly QC was conducted using *Blobtools* (Laetsch & Blaxter, 2017). Pacbio contigs were scaffolded by the Tell-seq data using *ARCS/LINKS* (Yeo et al., 2017). *BBMap* (Bushnell, 2014) was used to check for containment in the resulting assemblies. Only two suspected bacterial sequences were identified and removed from the J1 assembly. Canonical insect telomeric repeats (TTAGG) were identified (see Supplementary Tables 1, 2). The final J1 assembly included 109 scaffolds with a total length of 239,102,137 bp, and an N50 scaffold size of 10,579,937 bp. J2 had 92 scaffolds with a total of 236,044,553 bp, and an N50 of 10,965,113 bp.

### Synteny analysis

To compare the genomes for J1 and J2, synteny plots (“*MUMmerplot*”) were created using *MUMmer* (Kurtz et al., 2004) based on whole-genome alignments between the J1 and J2 assemblies, as well as each aligned against the *Pogonomyrmex californicus* genome (ver. 3.1, https://www.ncbi.nlm.nih.gov/datasets/genome/GCA_050084945.1/), which is the only available genome from the *Pogonomyrmex* genus that is assembled to chromosome level. The *nucmer* program of *MUMmer* was applied to align the genomes and the alignments were filtered using the *delta-filter* program. Finally, the *mummerplot* program was used to produce the synteny plots. For a closer inspection of the scaffold 1 gap region (8,146,955-13,350,929bp) we repeated the analysis described above only with that segment of the J1 chromosome. Orthologous genes were used as an alternative approach for synteny analysis. Predicted protein-coding sequences from *P. californicus* were blasted (*blastn*) against both reference genomes of J1 and J2 (Camacho et al., 2009). The synteny plot was based only on the genes that had a unique hit in each of the genomes with a 1e-150 e-value cutoff.

### Whole genome sequencing of males

Twelve males of each lineage were whole-genome sequenced on an Illumina HiSeq 3000X sequencer using 150 paired-end sequencing. Resulting reads were quality-checked using *fastQC*. First, *fastuniq* (Xu et al., 2012) removed read duplicates that occurred during Illumina sequencing. Then, reads were trimmed using *TRIMMOMATIC* with the following parameters: TruSeq3-PE-2.fa:2:30:10 SLIDINGWINDOW:4:15 MINLEN:75 (Bolger et al., 2014) and overlapping paired-read short reads were combined using *FLASH* (Magoč et al., 2011). *bowtie2* with the --end-to-end and --very-sensitive options was used to align the reads of all 24 samples to both J1 and J2 reference genomes. Reads mapped by *bowtie2* were filtered for high alignment score (alignment score AS > -25, i.e., allowing up to 4 mismatching bases) and unique mapping (second best alignment score XS < -50). Average coverage of 10-18X per sample was achieved. The genome coverage was plotted for each sample using *samtools depth* (Danecek et al., 2021), with coverage averaged over 1 Mb windows. The HaplotypeCaller of the *Genome Analysis Toolkit (GATK)* (McKenna et al., 2010) was used to call a total of 1,070,494 SNPs.

### Genomic (ddRAD) sequencing of workers

The worker samples were analyzed using reduced representation genomic sequencing - the double-digest restriction site associated DNA (ddRAD) sequencing protocol. For the full details please see Glinka et al. (2026). Briefly, DNA was extracted from the head and thorax of each worker sample using the Qiagen DNeasy blood and tissue protocol followed by ethanol precipitation. We used the ddRAD sequencing protocol following Brelsford et al. (2016), which was modified from the original protocol by Parchman et al. (2012). Libraries were sequenced by 150b paired-end Illumina sequencing. The resulting sequencing reads were processed with the *Stacks* v. 2 pipeline (Catchen et al., 2011). Briefly, reads were mapped to the *P. barbatus* reference genome of the J1 lineage using *bowtie2* v. 2.5.3 (Langmead & Salzberg, 2012). The *ref_map.pl* program was used to call SNPs. The SNPs were filtered for the number of missing genotypes (--max-missing 0.75), depth (--min-DP 6), and minor allele frequency (--maf 0.01) using *VCFtools* (Danecek et al., 2011).

The resulting genotype data from 1101 worker samples in 25,677 SNPs were used to calculate the percentage of heterozygous genotypes in each SNPusing *VCFtools* (Danecek et al., 2011).

### Genome annotation

We used *GeMoMa* (version 1.9; Keilwagen et al., 2016, 2018) to annotate the genes of *P. barbatus* J1-using *ab initio* gene prediction with *Augustus* (Stanke et al., 2006) and the following insect species as references for gene homology (GenBank references in brackets): *Pogonomyrmex barbatus* (GCF_000187915.1), *P. californicus* (GCA_050084945.1), *Camponotus floridanus* (GCF_003227725.1), *Harpegnathos saltator* (GCF_003227715.1), *Linepithema humile* (GCF_000217595.1), *Solenopsis invicta* (GCF_016802725.1), *Formica exsecta* (GCF_003651465.1), *Anopheles gambiae* (GCF_000005575.2), *Apis mellifera* (GCF_003254395.2), *Drosophila melanogaster* (GCF_000001215.4), *Nasonia vitripennis* (GCF_009193385.2), *Tribolium castaneum* (GCF_000002335.3). Then we blasted (*blastp*) (Camacho et al., 2009) the predicted protein sequence for the longest transcript of each gene against the Swissprot database (Bateman et al., 2025) and transferred the functional annotation from the top hit to the *P. barbatus* gene, including gene description and GO terms. Additionally, we used the *Happy-ABCENTH* pipeline (version 1.0; https://github.com/biorover/HAPpy-ABCENTH) to annotate the ORs in the genome. We then replaced any gene annotations by *GeMoMa* with overlapping annotations from *Happy-ABCENTH*. We conducted enrichment tests for the two gene sets (J1 scaffold 1 and the synteny gap) using Fisher’s exact test as implemented in the *topGO* package (Alexa & Rahnenführer, 2025). P-values were corrected using the Benjamini-Hochberg correction (Benjamini & Hochberg, 1995).

### Odorant receptor gene tree

We constructed a gene tree for OR subfamily L to investigate the evolutionary history of this gene cluster in J1 scaffold 1. We used predicted OR protein sequences (Vizueta et al., 2025) for eight representative species from diverged ant subfamilies: *Pogonomyrmex barbatus* (Myrmicinae), *Myrmica rubra* (Myrmicinae), *Solenopsis invicta* (Myrmicinae), *Camponotus fellah* (Formicinae), *Linepithema humile* (Dolichoderinae), *Harpegnathos saltator* (Ponerinae), *Proceratium itoi* (Proceratiinae), and *Leptanilla species* (Leptanillinae). We aligned the sequences using MAFFT version 7.511 (Katoh & Standley, 2013) with the L-ins-i algorithm. We reconstructed the gene tree using RAxML version 8.0.26 (Stamatakis, 2014) with the PROTCATLG model and 100 bootstrap repeats. The tree was visualized using Figtree version 1.4.4 (Rambaut, 2018).

## Acknowledgements

The work was supported by the U.S.-Israel Binational Science Foundation grant (2019655) to E.P. and National Science Foundation Award (1940647) to D.M.G. We are grateful to Aparna Lajmi and Chih-Chi Lee for guidance in the lab. Further, we would like to thank Alvaro Gonzalo Hernandez, Christopher J Fields, Gloria A Rendon, Chris Wright, and Elizabeth Kaitlin Hogan for their support with Tell-seq and PacBio sequencing.

## Authors’ contributions

F.G.: methodology, investigation, formal analysis, data curation, writing – original draft, visualization

Y.P.: writing – original draft, visualization Z.F.: visualization

K.W.: data curation

D.M.G.: resources, supervision, project administration, writing – original draft, funding acquisition

E.P.: conceptualization, methodology, resources, supervision, project administration, writing – original draft, funding acquisition

## Data Accessibility Statement

The reference genomes were deposited at NCBI GenBank under the accession JBVVXZ000000000 and JBVVYA000000000. The version described in this paper is version JBVVXZ010000000 and JBVVYA010000000. The 24 male WGS samples are being processed to NCBI (PRJNA1434043). Gene annotations and predicted protein sequences were submitted to the Zenodo repository 10.5281/zenodo.20110933.

## Competing Interests Statement

The authors declare no conflicts of interest.

## Supplementary

**Supplementary Figure 1:**
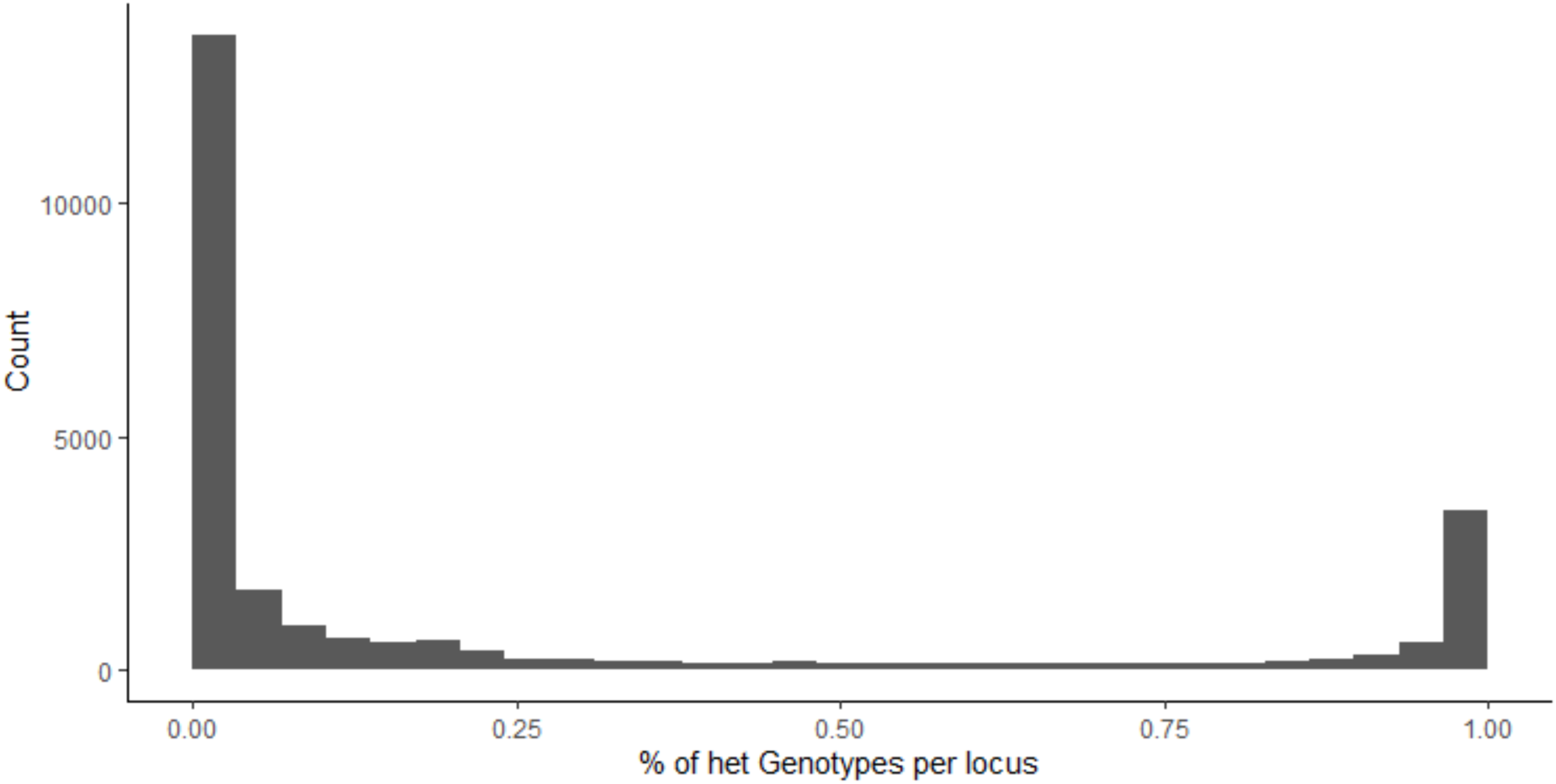
Distribution of allele frequency per single nucleotide polymorphism locus in the worker samples. The U-shaped distribution is in line with the expectation for F1 hybrids between genetically diverged populations.

**Supplementary Figure 2:**
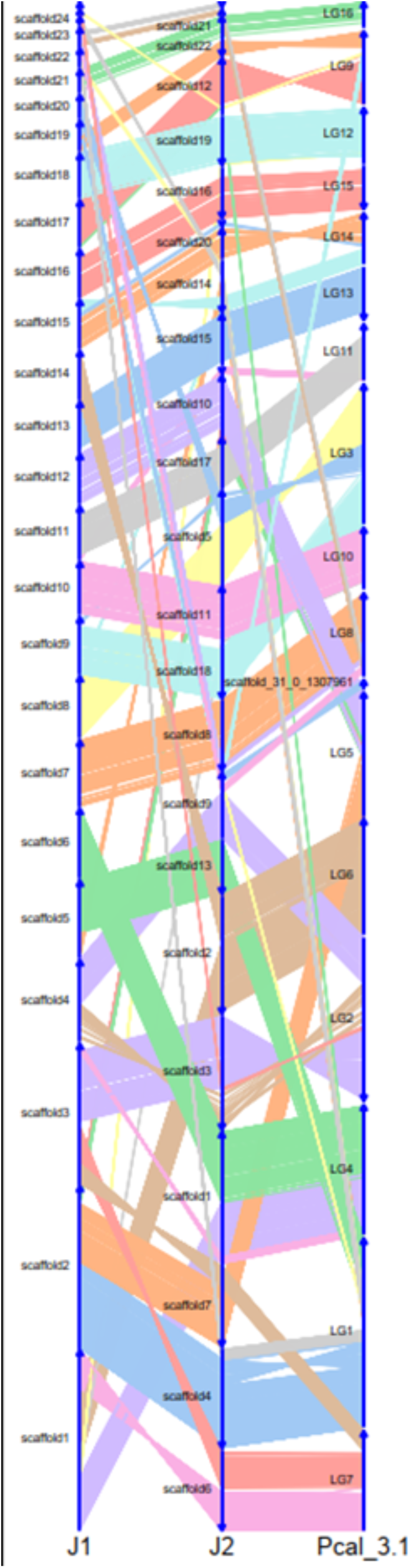
Conservation of synteny illustrated using a river plot comparing the J1 and J2 of *P. barbatus* and *P. californicus* genomes. Each line represents one-to-one orthologous protein-coding genes.

**Supplementary Figure 3:**
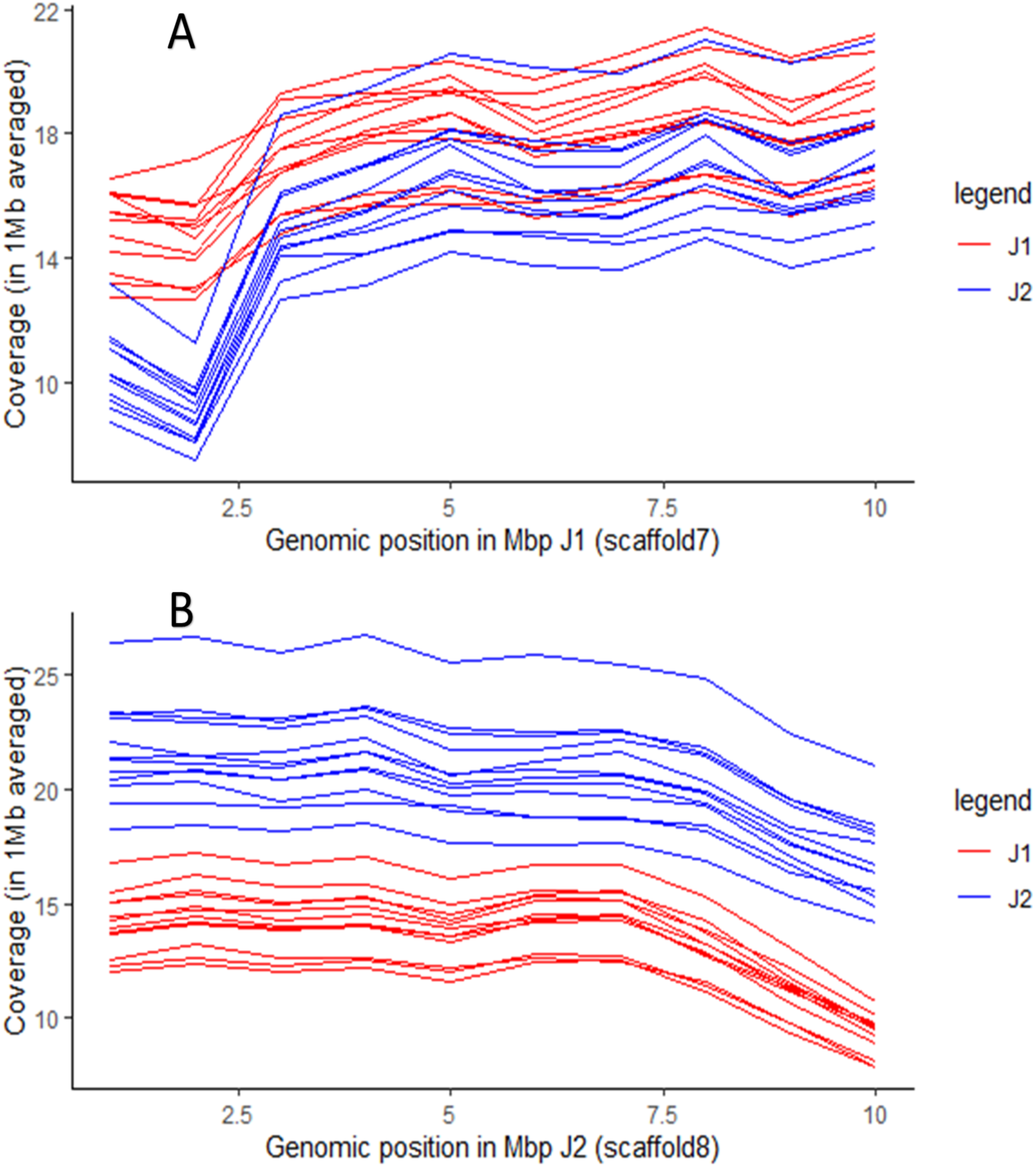
Coverage plots for male samples. Red: J1 samples, blue: J2 samples. **A:** scaffold 7 of J1; **B:** scaffold 8 of J2, which is the homologous scaffold.

**Supplementary Figure 4:**
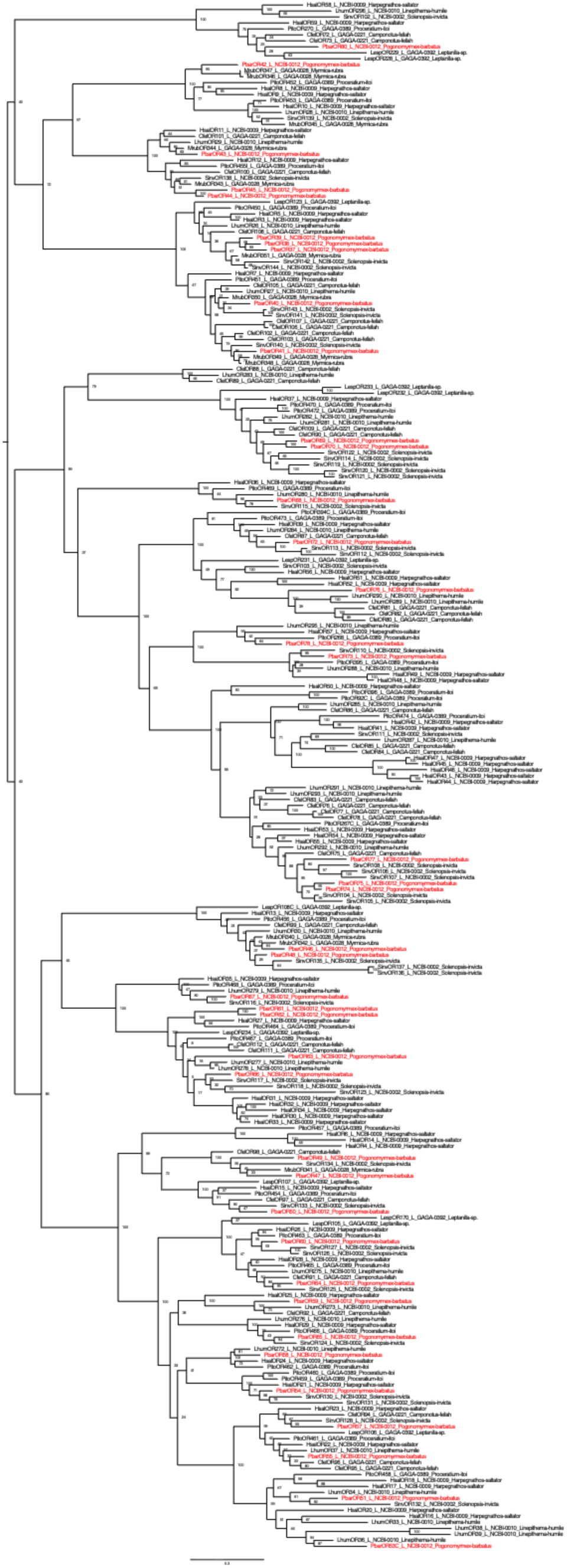
Gene tree of subfamily L of the odorant receptor (OR) gene family. The tree includes subfamily L ORs from eight representative species of diverged ant subfamilies. *P. barbatus* genes are colored in red.

**Supplementary Table 1:** List of the large scaffolds (>70,000 bp) for the J1 reference genome. Telomeric repeats indicated where identified.

| Scaffold name | Length | Left side telomere repeat | Right side telomere repeat |
| --- | --- | --- | --- |
| scaffold1 | 29,218,669 |  | TTAAGGTTAGG |
| scaffold2 | 26,152,519 | TAACCTAACC |  |
| scaffold3 | 22,607,820 | TAACCTAACC | TTAAGGTTAGG |
| scaffold4 | 13,109,157 |  | TTAAGGTTAGG |
| scaffold5 | 12,417,964 |  | TTAAGGTTAGG |
| scaffold6 | 11,203,750 |  | TTAAGGTTAGG |
| scaffold7 | 10,579,937 |  | TTAAGGTTAGG |
| scaffold8 | 9,943,993 | TAACCTAACC | TTAAGGTTAGG |
| scaffold9 | 9,097,628 | TAACCTAACC |  |
| scaffold10 | 8,521,807 |  | TTAAGGTTAGG |
| scaffold11 | 8,501,584 | TAACCTAACC |  |
| scaffold12 | 8,028,656 |  | TTAAGGTTAGG |
| scaffold13 | 7,989,518 | TAACCTAACC | TTAAGGTTAGG |
| scaffold14 | 7,785,685 | TAACCTAACC |  |
| scaffold15 | 7,774,943 | TAACCTAACC |  |
| scaffold16 | 7,642,452 | TAACCTAACC |  |
| scaffold17 | 7,502,175 |  | TTAAGGTTAGG |
| scaffold18 | 7,088,262 |  | TTAAGGTTAGG |
| scaffold19 | 4,868,040 | TAACCTAACC |  |
| scaffold20 | 3,730,139 |  |  |
| scaffold21 | 3,581,242 | TAACCTAACC | TTAAGGTTAGG |
| scaffold22 | 2,959,188 |  |  |
| scaffold23 | 2,903,150 | TAACCTAACC |  |
| scaffold24 | 1,294,252 |  |  |
| scaffold25 | 722,980 |  |  |
| scaffold26 | 642,951 |  |  |
| scaffold27 | 434,911 | TAACCTAACC |  |
| scaffold28 | 429,889 | TAACCTAACC |  |
| scaffold29 | 309,341 | TAACCTAACC |  |
| scaffold30 | 226,601 | TAACCTAACC |  |
| scaffold31 | 191,036 | TAACCTAACC |  |

**Supplementary Table 2:** List of the large scaffolds (>70,000 bp) for the J2 reference genome. Telomeric repeats indicated where identified..

| Scaffold name | Length | Left side Telomere Repeat | Right Side Telomere Repeat |
| --- | --- | --- | --- |
| scaffold1 | 20,770,626 | TAACCTAACC |  |
| scaffold2 | 18,567,161 | TAACCTAACC | TTAAGGTTAGG |
| scaffold3 | 17,919,464 | TAACCTAACC |  |
| scaffold4 | 15,619,943 |  | TTAAGGTTAGG |
| scaffold5 | 14,732,959 | TAACCTAACC |  |
| scaffold6 | 12,627,774 |  | TAACCTAACC |
| scaffold7 | 12,626,827 | TAACCTAACC | TTAAGGTTAGG |
| scaffold8 | 10,965,113 | TAACCTAACC |  |
| scaffold9 | 10,197,547 |  |  |
| scaffold10 | 9,347,904 | TTAAGGTTAGG |  |
| scaffold11 | 9,283,530 | TAACCTAACC | TTAAGGTTAGG |
| scaffold12 | 8,872,895 |  | TTAAGGTTAGG |
| scaffold13 | 8,347,315 | TAACCTAACC |  |
| scaffold14 | 8,103,180 |  | TTAAGGTTAGG |
| scaffold15 | 7,961,234 |  |  |
| scaffold16 | 7,924,686 | TAACCTAACC | TTAAGGTTAGG |
| scaffold17 | 7,910,573 |  |  |
| scaffold18 | 7,848,957 |  | TTAAGGTTAGG |
| scaffold19 | 7,444,109 | TAACCTAACC |  |
| scaffold20 | 4,521,639 | TAACCTAACC |  |
| scaffold21 | 3,249,046 |  | TTAAGGTTAGG |
| scaffold22 | 2,427,047 |  | TTAAGGTTAGG |
| scaffold23 | 1,138,895 |  | TTAAGGTTAGG |
| scaffold24 | 957,398 |  |  |
| scaffold25 | 856,613 |  |  |
| scaffold26 | 820,652 | TAACCTAACC |  |
| scaffold27 | 793,817 |  | TTAAGGTTAGG |
| scaffold28 | 696,027 |  |  |
| scaffold29 | 604,743 | TAACCTAACC |  |
| scaffold30 | 500,787 |  | TTAAGGTTAGG |
| scaffold31 | 479,463 |  | TTAAGGTTAGG |
| scaffold32 | 421,527 |  |  |
| scaffold33 | 244,862 |  |  |
| scaffold34 | 211,420 |  |  |
| scaffold35 | 70,598 | TAACCTAACC |  |

**Supplementary Table 3:** Gene counts for OR subfamilies in J1 scaffold 1.

| Subfamily | Count | Total number in genome |
| --- | --- | --- |
| K | 2 | 2 |
| U | 5 | 26 |
| L | 43 | 43 |
| N | 3 | 3 |
| P | 8 | 8 |
| M | 4 | 4 |
| J | 2 | 2 |
| 9E (9-exon) | 51 | 152 |

## Notes

https://doi.org/10.5281/zenodo.20110933

### Competing Interest Statement

The authors have declared no competing interest.

## Bibliography

Alexa, A., & Rahnenführer, J. (2025). topGO: Enrichment Analysis for Gene Ontology. doi:10.18129/B9.bioc.topGOdoi:10.18129/B9.bioc.topGO, R package version 2.62.0, https://bioconductor.org/packages/topGO

Anderson, K. E., Gadau, J., Mott, B. M., Johnson, R. A., Altamirano, A., Strehl, C., & Fewell, J. H. (2006). Distribution and evolution of genetic caste determination in Pogonomyrmex seed-harvester ants. Ecology, 87(9), 2171–2184. 10.1890/0012-9658(2006)87

Anderson, K. E., Linksvayer, T. A., & Smith, C. R. (2008). The causes and consequences of genetic caste determination in ants (Hymenoptera: Formicidae). Myrmecological News, 11, 119--132.

Bateman, A., Martin, M. J., Orchard, S., Magrane, M., Adesina, A., Ahmad, S., Bowler-Barnett, E. H., Bye-A-Jee, H., Carpentier, D., Denny, P., Fan, J., Garmiri, P., da Costa Gonzales, L. J., Hussein, A., Ignatchenko, A., Insana, G., Ishtiaq, R., Joshi, V., Jyothi, D., … Zhang, J. (2025). UniProt: the Universal Protein Knowledgebase in 2025. Nucleic Acids Research, 53(D1), D609–D617. 10.1093/NAR/GKAE1010

Benirschke, K., Brownhill, L. E., & Beath, M. M. (1962). Somatic chromosomes of the horse, the donkey and their hybrids, the mule and the hinny. Journal of Reproduction and Fertility, 4(3), 319–326. 10.1530/jrf.0.0040319

Benjamini, Y., & Hochberg, Y. (1995). Controlling the False Discovery Rate: A Practical and Powerful Approach to Multiple Testing. Journal of the Royal Statistical Society: Series B (Methodological), 57(1), 289–300. 10.1111/J.2517-6161.1995.TB02031.X

Beukeboom, L. W., & Perrin, N. (2014). The evolution of sex chromosomes. The Evolution of Sex Determination, 89–114. 10.1093/acprof:oso/9780199657148.003.0005

Blackmon, H., & Demuth, J. P. (2015). Genomic origins of insect sex chromosomes. Current Opinion in Insect Science, 7, 45–50. 10.1016/J.COIS.2014.12.003

Blomquist, G. J., & Bagnères, A. G. (2010). Insect hydrocarbons biology, biochemistry, and chemical ecology. *Insect Hydrocarbons Biology*, Biochemistry, and Chemical Ecology, 1–492. 10.1017/CBO9780511711909

Bolger, A. M., Lohse, M., & Usadel, B. (2014). Trimmomatic: a flexible trimmer for Illumina sequence data. Bioinformatics, 30(15), 2114–2120. 10.1093/BIOINFORMATICS/BTU170

Brelsford, A., Dufresnes, C., & Perrin, N. (2016). High-density sex-specific linkage maps of a European tree frog (Hyla arborea) identify the sex chromosome without information on offspring sex. Heredity, 116(2), 177–181. 10.1038/hdy.2015.83

Bushnell, B. (2014). BBMap: A Fast, Accurate, Splice-Aware Aligner.

Camacho, C., Coulouris, G., Avagyan, V., Ma, N., Papadopoulos, J., Bealer, K., & Madden, T. L. (2009). BLAST+: architecture and applications. BMC Bioinformatics, 10. 10.1186/1471-2105-10-421

Castillo, E. R., Marti, D. A., & Bidau, C. J. (2010). Sex and Neo-Sex Chromosomes in Orthoptera: A Review. Journal of Orthoptera Research, 19(2), 213–231. 10.1665/034.019.0207

Catchen, J. M., Amores, A., Hohenlohe, P., Cresko, W., & Postlethwait, J. H. (2011). Stacks: Building and genotyping loci de novo from short-read sequences. G3: Genes, Genomes, Genetics, 1(3), 171–182. 10.1534/g3.111.000240

Charlesworth, D. (2017). Evolution of recombination rates between sex chromosomes. Philosophical Transactions of the Royal Society B: Biological Sciences, 372(1736). 10.1098/rstb.2016.0456

Cheng, H., Concepcion, G. T., Feng, X., Zhang, H., & Li, H. (2021). Haplotype-resolved de novo assembly using phased assembly graphs with hifiasm. Nature Methods, 18(2), 170–175. 10.1038/S41592-020-01056-5

Danecek, P., Auton, A., Abecasis, G., Albers, C. A., Banks, E., DePristo, M. A., Handsaker, R. E., Lunter, G., Marth, G. T., Sherry, S. T., McVean, G., & Durbin, R. (2011). The variant call format and VCFtools. Bioinformatics, 27(15), 2156–2158. 10.1093/BIOINFORMATICS/BTR330

Danecek, P., Bonfield, J. K., Liddle, J., Marshall, J., Ohan, V., Pollard, M. O., Whitwham, A., Keane, T., McCarthy, S. A., & Davies, R. M. (2021). Twelve years of SAMtools and BCFtools. GigaScience, 10(2), 1–4. 10.1093/GIGASCIENCE/GIAB008

Darras, H., Araujo, N. D. S., Baudry, L., Guiglielmoni, N., Lorite, P., Marbouty, M., Rodriguez, F., Arkhipova, I., Koszul, R., Flot, J. F., & Aron, S. (2022). Chromosome-level genome assembly and annotation of two lineages of the ant Cataglyphis hispanica: stepping stones towards genomic studies of hybridogenesis and thermal adaptation in desert ants. Peer Community Journal, 2. 10.24072/PCJOURNAL.140/

Dimitri, P., & Junakovic, N. (1999). Revising the selfish DNA hypothesis: new evidence on accumulation of transposable elements in heterochromatin. Trends in Genetics, 15(4), 123–124.

Eliyahu, D., Ross, K. G., Haight, K. L., Keller, L., & Liebig, J. (2011). Venom Alkaloid and Cuticular Hydrocarbon Profiles Are Associated with Social Organization, Queen Fertility Status, and Queen Genotype in the Fire Ant Solenopsis invicta. Journal of Chemical Ecology, 37(11), 1242–1254. 10.1007/s10886-011-0037-y

Ezaz, T., Sarre, S. D., O’Meally, D., Marshall Graves, J. A., & Georges, A. (2009). Sex Chromosome Evolution in Lizards: Independent Origins and Rapid Transitions. Cytogenetic and Genome Research, 127(2–4), 249–260. 10.1159/000300507

Fishman, L., Stathos, A., Beardsley, P. M., Williams, C. F., & Hill, J. P. (2013). Chromosomal Rearrangements and the Genetics of Reproductive Barriers in Mimulus (Monkey Flowers). Evolution, 67(9), 2547–2560. 10.1111/EVO.12154

Fontana, S., Chang, N. C., Chang, T., Lee, C. C., Dang, V. D., & Wang, J. (2020). The fire ant social supergene is characterized by extensive gene and transposable element copy number variation. Molecular Ecology, 29(1), 105–120.

Glinka, F., Steiner, E. B., Privman, E., & Gordon, D. M. (2026). Shifts in demography in changing ecological conditions in a dependent-lineage population of harvester ant colonies. BioRxiv. 10.64898/2026.03.06.710041

Gordon, D. M. (1995). The development of an ant colony’s foraging range. Animal Behaviour, 49(3), 649–659. 10.1016/0003-3472(95)80198-7

Gotzek, D., Robertson, H. M., Wurm, Y., & Shoemaker, D. W. (2011). Odorant Binding Proteins of the Red Imported Fire Ant, Solenopsis invicta: An Example of the Problems Facing the Analysis of Widely Divergent Proteins. PLOS ONE, 6(1), e16289. 10.1371/journal.pone.0016289

Graber, L. C., Johnson, R. A., & Moreau, C. S. (2024). UCE phylogenomics inform the systematics and geographic range evolution of the harvester ant genus Pogonomyrmex. BioRxiv, 2024.11.12.623263. 10.1101/2024.11.12.623263

Greene, M. J., & Gordon, D. M. (2003). Cuticular hydrocarbons inform task decisions. Nature, 423(6935), 32–32. 10.1038/423032a

Helms Cahan, S., Goodman, P., & Grauer, J. A. (2023). Ecological and genetic distinctiveness of socially hybridogenetic lineages of Pogonomyrmex harvester ants at regional and local scales. Evolutionary Ecology, 37(4), 645–667. 10.1007/S10682-023-10241-9

Helms Cahan, S., & Keller, L. (2003). Complex hybrid origin of genetic caste determination in harvester ants. Nature, 424(6946), 306–309. 10.1038/nature01744

Helms Cahan, S., Nguyen, A. D., & Zhou, Y. (2022). Population genomics supports multiple hybrid zone origins of socially hybridogenetic lineages of Pogonomyrmex harvester ants. Evolution, 76(5), 1016–1032. 10.1111/EVO.14481

Helms Cahan, S., Parker, J. D., Rissing, S. W., Johnson, R. A., Polony, T. S., Weiser, M. D., & Smith, D. R. (2002). Extreme genetic differences between queens and workers in hybridizing Pogonomyrmex harvester ants. Proceedings of the Royal Society of London. Series B: Biological Sciences, 269(1503), 1871–1877. 10.1098/RSPB.2002.2061

Julian, G. E., Fewell, J. H., Gadau, J., Johnson, R. A., & Larrabee, D. (2002). Genetic determination of the queen caste in an ant hybrid zone. Proceedings of the National Academy of Sciences, 99(12), 8157–8160. 10.1073/pnas.112222099

Kaiser, V. B., & Bachtrog, D. (2010). Evolution of sex chromosomes in insects. Annual Review of Genetics, 44(Volume 44, 2010), 91–112. 10.1146/ANNUREV-GENET-102209-163600/CITE/REFWORKS

Katoh, K., & Standley, D. M. (2013). MAFFT Multiple Sequence Alignment Software Version 7: Improvements in Performance and Usability. Molecular Biology and Evolution, 30(4), 772–780. 10.1093/molbev/mst010

Keilwagen, J., Hartung, F., Paulini, M., Twardziok, S. O., & Grau, J. (2018). Combining RNA-seq data and homology-based gene prediction for plants, animals and fungi. BMC Bioinformatics, 19(1). 10.1186/S12859-018-2203-5

Keilwagen, J., Wenk, M., Erickson, J. L., Schattat, M. H., Grau, J., & Hartung, F. (2016). Using intron position conservation for homology-based gene prediction. Nucleic Acids Research, 44(9), e89–e89. 10.1093/NAR/GKW092

Keller, L., & Ross, K. G. (1998). Selfish genes: a green beard in the red fire ant. Nature 1998 394:6693, 394(6693), 573–575. 10.1038/29064

Krieger, M. J. B., & Ross, K. G. (2002). Identification of a major gene regulating complex social behavior. Science (New York, N.Y.), 295(5553), 328–332. 10.1126/science.1065247

Kurtz, S., Phillippy, A., Delcher, A. L., Smoot, M., Shumway, M., Antonescu, C., & Salzberg, S. L. (2004). Open Access Versatile and open software for comparing large genomes. 5(2), 12. http://www.tigr.org/software/mummer.

Laetsch, D. R., & Blaxter, M. L. (2017). BlobTools: Interrogation of genome assemblies. F1000Research 2017 6:1287, *6*, 1287. 10.12688/f1000research.12232.1

Langmead, B., & Salzberg, S. L. (2012). Fast gapped-read alignment with Bowtie 2. Nature Methods, 9(4), 357–359. 10.1038/nmeth.1923

Linksvayer, T. A. (2006). Direct, maternal, and sibsocial genetic effects on individual and colony traits in an ant. Evolution, 60(12), 2552–2561. 10.1111/J.0014-3820.2006.TB01889.X

Magoč, T., Magoč, M., & Salzberg, S. L. (2011). FLASH: fast length adjustment of short reads to improve genome assemblies. 27(21), 2957–2963. 10.1093/bioinformatics/btr507

McKenna, A., Hanna, M., Banks, E., Sivachenko, A., Cibulskis, K., Kernytsky, A., Garimella, K., Altshuler, D., Gabriel, S., Daly, M., & DePristo, M. A. (2010). The Genome Analysis Toolkit: A MapReduce framework for analyzing next-generation DNA sequencing data. Genome Research, 20(9), 1297–1303. 10.1101/GR.107524.110

McKenzie, S. K., Fetter-Pruneda, I., Ruta, V., & Kronauer, D. J. C. (2016). Transcriptomics and neuroanatomy of the clonal raider ant implicate an expanded clade of odorant receptors in chemical communication. Proceedings of the National Academy of Sciences, 113(49), 14091–14096. 10.1073/pnas.1610800113

Meisel, R. P., Olafson, P. U., Adhikari, K., Guerrero, F. D., Konganti, K., & Benoit, J. B. (2020). Sex Chromosome Evolution in Muscid Flies. G3 Genes|Genomes|Genetics, 10(4), 1341–1352. 10.1534/G3.119.400923

Mott, B. M., Gadau, J., Anderson, K. E., Correspondence, B. M., Mott, U.-A., & Carl, H. (2015). Phylogeography of Pogonomyrmex barbatus and P. rugosus harvester ants with genetic and environmental caste determination. Ecology and Evolution, 5(14), 2798–2826. 10.1002/ECE3.1507

Ohno, S. (1967) Sex chromosomes and sex-linked genes. Springer, New York.

Parchman, T. L., Gompert, Z., Mudge, J., Schilkey, F. D., Benkman, C. W., & Buerkle, C. A. (2012). Genome-wide association genetics of an adaptive trait in lodgepole pine. Molecular Ecology, 21(12), 2991–3005. 10.1111/j.1365-294X.2012.05513.x

Pask, G. M., Slone, J. D., Millar, J. G., Das, P., Moreira, J. A., Zhou, X., Bello, J., Berger, S. L., Bonasio, R., Desplan, C., Reinberg, D., Liebig, J., Zwiebel, L. J., & Ray, A. (2017). Specialized odorant receptors in social insects that detect cuticular hydrocarbon cues and candidate pheromones. Nature Communications 2017 8:1, 8(1), 297-. 10.1038/s41467-017-00099-1

Pennell, M. W., Kirkpatrick, M., Otto, S. P., Vamosi, J. C., Peichel, C. L., Valenzuela, N., & Kitano, J. (2015). Y Fuse? Sex Chromosome Fusions in Fishes and Reptiles. PLOS Genetics, 11(5), e1005237. 10.1371/journal.pgen.1005237

Rambaut, A. (2018). FigTree v1.4.4. https://tree.bio.ed.ac.uk/software/figtree/

Ranz, J. M., Casals, F., & Ruiz, A. (2001). How Malleable is the Eukaryotic Genome? Extreme Rate of Chromosomal Rearrangement in the Genus Drosophila. Genome Research, 11(2), 230–239. 10.1101/gr.162901

Rieseberg, L. H. (2001). Chromosomal rearrangements and speciation. Trends in Ecology and Evolution, 16(7), 351–358. 10.1016/s0169-5347(01)02187-5

Romiguier, J., Cameron, S. A., Woodard, S. H., Fischman, B. J., Keller, L., & Praz, C. J. (2016). Phylogenomics Controlling for Base Compositional Bias Reveals a Single Origin of Eusociality in Corbiculate Bees. Molecular Biology and Evolution, 33(3), 670–678. 10.1093/MOLBEV/MSV258

Romiguier, J., Fournier, A., Yek, S. H., & Keller, L. (2017). Convergent evolution of social hybridogenesis in Messor harvester ants. Molecular Ecology, 26(4), 1108–1117. 10.1111/MEC.13899

Schwander, T., Helms Cahan, S., & Keller, L. (2007). Characterization and distribution of Pogonomyrmex harvester ant lineages with genetic caste determination. Molecular Ecology, 16(2), 367–387. 10.1111/j.1365-294X.2006.03124.x

Schwander, T., Libbrecht, R., & Keller, L. (2014). Supergenes and complex phenotypes. Current Biology, 24(7), R288–R294. 10.1016/j.cub.2014.01.056

Sirviö, A., Pamilo, P., Johnson, R. A., Page, R. E., & Gadau, J. (2011). Origin and Evolution of the Dependent Lineages in the Genetic Caste Determination System of Pogonomyrmex Ants. Evolution, 65(3), 869–884. 10.1111/J.1558-5646.2010.01170.X

Slone, J. D., Pask, G. M., Ferguson, S. T., Millar, J. G., Berger, S. L., Reinberg, D., Liebig, J., Ray, A., & Zwiebel, L. J. (2017). Functional characterization of odorant receptors in the ponerine ant, Harpegnathos saltator. Proceedings of the National Academy of Sciences of the United States of America, 114(32), 8586–8591. 10.1073/pnas.1704647114

Smith, C. D., Zimin, A., Holt, C., Abouheif, E., Benton, R., Cash, E., Croset, V., Currie, C. R., Elhaik, E., Elsik, C. G., Fave, M. J., Fernandes, V., Gadau, J., Gibson, J. D., Graur, D., Grubbs, K. J., Hagen, D. E., Helmkampf, M., Holley, J. A., … Tsutsui, N. D. (2011a). Draft genome of the globally widespread and invasive Argentine ant (Linepithema humile). Proceedings of the National Academy of Sciences, 108(14), 5673–5678. 10.1073/pnas.1008617108

Smith, C. R., Smith, C. D., Robertson, H. M., Helmkampf, M., Zimin, A., Yandell, M., Holt, C., Hu, H., Abouheif, E., Benton, R., Cash, E., Croset, V., Currie, C. R., Elhaik, E., Elsik, C. G., Favé, M. J., Fernandes, V., Gibson, J. D., Graur, D., … Gadau, J. (2011b). Draft genome of the red harvester ant Pogonomyrmex barbatus. Proceedings of the National Academy of Sciences of the United States of America, 108(14), 5667–5672.

Sprenger, P. P., & Menzel, F. (2020). Cuticular hydrocarbons in ants (Hymenoptera: Formicidae) and other insects: how and why they differ among individuals, colonies, and species. Myrmecological News, 30, 31–46. 10.25849/myrmecol.news

Stamatakis, A. (2014). RAxML version 8: a tool for phylogenetic analysis and post-analysis of large phylogenies. Bioinformatics, 30(9), 1312–1313. 10.1093/BIOINFORMATICS/BTU033

Stanke, M., Keller, O., Gunduz, I., Hayes, A., Waack, S., & Morgenstern, B. (2006). AUGUSTUS: ab initio prediction of alternative transcripts. Nucleic Acids Research, 34(Web Server issue). 10.1093/NAR/GKL200

Steinemann, S., & Steinemann, M. (2005). Y chromosomes: Born to be destroyed. BioEssays, 27(10), 1076–1083. 10.1002/bies.20288

Sundaram, M., Steiner, E., & Gordon, D. M. (2022). Rainfall, neighbors, and foraging: The dynamics of a population of red harvester ant colonies 1988–2019. Ecological Monographs, 92(2), e1503. 10.1002/ECM.1503

Traut, W. (2010). New Y chromosomes and early stages of sex chromosome differentiation: Sex determination in Megaselia. Journal of Genetics, 89(3), 307–313. 10.1007/S12041-010-0042-X/METRICS

Uno, Y., Nishida, C., Oshima, Y., Yokoyama, S., Miura, I., Matsuda, Y., & Nakamura, M. (2008). Comparative chromosome mapping of sex-linked genes and identification of sex chromosomal rearrangements in the Japanese wrinkled frog (*Rana rugosa*, Ranidae) with ZW and XY sex chromosome systems. Chromosome Research, 16(4), 637–647. 10.1007/S10577-008-1217-7/METRICS

Vizueta, J., Xiong, Z., Ding, G., Larsen, R. S., Ran, H., Gao, Q., … & Zhang, G. (2025). Adaptive radiation and social evolution of the ants. Cell, 188(18), 4828–4848. 10.1016/j.cell.2025.05.030

Volny, V. P., & Gordon, D. M. (2002). Genetic basis for queen-worker dimorphism in a social insect. Proceedings of the National Academy of Sciences of the United States of America, 99(9), 6108–6111. 10.1073/pnas.092066699

Volny, V. P., Greene, M. J., & Gordon, D. M. (2006). Brood production and lineage discrimination in the red harvester ant (Pogonomyrmex barbatus). Ecology, 87(9), 2194–2200. 10.1890/0012-9658(2006)87[2194:BPALDI]2.0.CO;2

Wang, Jianping, Na, J. K., Yu, Q., Gschwend, A. R., Han, J., Zeng, F., Aryal, R., VanBuren, R., Murray, J. E., Zhang, W., Navajas-Pérez, R., Feltus, F. A., Lemke, C., Tong, E. J., Chen, C., Wai, C. M., Singh, R., Wang, M. L., Min, X. J., … Ming, R. (2012). Sequencing papaya X and Y h chromosomes reveals molecular basis of incipient sex chromosome evolution. Proceedings of the National Academy of Sciences of the United States of America, 109(34), 13710–13715. 10.1073/pnas.1207833109

Wang, J., Wurm, Y., Nipitwattanaphon, M., Riba-Grognuz, O., Huang, Y. C., Shoemaker, D., & Keller, L. (2013). A Y-like social chromosome causes alternative colony organization in fire ants. Nature, 493(7434), 664–668. 10.1038/nature11832

Weaver, N. (1955). Rearing of Honeybee Larvae on Royal Jelly in the Laboratory. *Science (New York*, N.Y.), 121(3145), 509–510. 10.1126/science.121.3145.509

Xu, H., Luo, X., Qian, J., Pang, X., Song, J., Qian, G., Chen, J., & Chen, S. (2012). FastUniq: A Fast De Novo Duplicates Removal Tool for Paired Short Reads. PLOS ONE, 7(12), e52249. 10.1371/JOURNAL.PONE.0052249

Yan, Z., Martin, S. H., Gotzek, D., Arsenault, S. V., Duchen, P., Helleu, Q., Riba-Grognuz, O., Hunt, B. G., Salamin, N., Shoemaker, D. W., Ross, K. G., & Keller, L. (2020). Evolution of a supergene that regulates a trans-species social polymorphism. Nature Ecology & Evolution 2020 4:2, 4(2), 240–249. 10.1038/s41559-019-1081-1

Yeo, S., Coombe, L., Chu, J., Warren, R. L., & Birol, I. (2017). ARCS: Assembly Roundup by Chromium Scaffolding. BioRxiv, 100750. 10.1101/100750

Zeng, H., Millar, J. G., Chen, L., Keller, L., & Ross, K. G. (2021). Characterization of Queen Supergene Pheromone in the Red Imported Fire Ant Using Worker Discrimination Assays. Journal of Chemical Ecology 2021 48:2, 48(2), 109–120. 10.1007/s10886-021-01336-0

